# Global chromosome topology and the two-component systems in concerted manner regulate transcription in *Streptomyces*

**DOI:** 10.1101/2021.09.15.460574

**Authors:** Martyna Gongerowska-Jac, Marcin J. Szafran, Jakub Mikołajczyk, Justyna Szymczak, Magda Bartyńska, Anna Gierlikowska, Sylwia Biały, Marie A. Elliot, Dagmara Jakimowicz

## Abstract

Bacterial gene expression is controlled at multiple levels, with chromosome supercoiling being one of the most global regulators. Global DNA supercoiling is maintained by the orchestrated action of topoisomerases. In *Streptomyces*, mycelial soil bacteria with a complex life cycle, topoisomerase I depletion led to elevated chromosome supercoiling, changed expression of significant fraction of genes, delayed growth and blocked sporulation. To identify supercoiling-induced sporulation regulators, we searched for *S. coelicolor* transposon mutants that were able to restore sporulation despite high chromosome supercoiling. We established that transposon insertion in genes encoding a novel two-component system named SatKR reversed the sporulation blockage resulting from topoisomerase I depletion. Transposition in *satKR* abolished the transcriptional induction of the genes within the so-called supercoiling-hypersensitive cluster (SHC). Moreover, we found that activated SatR also induced the same set of SHC genes under normal supercoiling conditions. We determined that the expression of genes in this region impacted *S. coelicolor* growth and sporulation. Interestingly, among the associated products is another two-component system (SitKR), indicating the potential for cascading regulatory effects driven by the SatKR and SitKR two-component systems. Thus, we demonstrated the concerted activity of chromosome supercoiling and a hierarchical two-component signalling system that impacts gene activity governing *Streptomyces* growth and sporulation.

**IMPORTANCE:** *Streptomyces*, soil bacteria with complex life cycle, are the producers of a broad range of biologically active compounds (e.g. antibiotics). *Streptomyces* respond to various environmental signals using complex transcriptional regulation mechanism. Understanding regulation of their gene expression is crucial for *Streptomyces* application as industrial organisms. Here, based on extensive transcriptomics analyses, we describe the concerted regulation of genes crucial for growth and development by global DNA supercoiling and novel two-component system. Our data indicate that regulated genes encode growth and sporulation regulator. Thus, we demonstrate that *Streptomyces* link the global regulatory strategies to adjusts life cycle to unfavourable conditions.

## INTRODUCTION

In all organisms, gene expression is precisely controlled, primarily at the level of transcription initiation. The main transcriptional regulatory factors include promoter DNA sequences and trans-acting transcriptional regulators. Bacterial genomes encode numerous transcriptional regulators, among which the key players are DNA-binding proteins like sigma factors, which recruit RNA polymerase (RNAP) to promoters, and transcription factors (TFs), which may act as repressors or activators by affecting RNAP binding (1, 2). Notably, the binding of regulatory proteins in bacteria is in turn controlled by systems that adjust transcription in response to external and internal cell conditions. Examples include changes in chromosome topology to modify promoter accessibility to regulatory factors, as well as the modification of regulatory protein activity itself (3).

The activity of specific sigma factors or regulatory proteins can be modulated through partner protein or ligand binding, as well as through proteolysis or covalent modifications like phosphorylation. The importance of phosphorylation has been well established for regulators that are a part of two-component systems (TCSs). Canonical TCSs consist of a transmembrane sensor histidine kinase (HK) and cytoplasmic response regulator (RR), which detects environmental signals and triggers intracellular responses, respectively (4). Upon signal sensing, the kinase in classical TCSs undergoes autophosphorylation and subsequently transfers the phosphate moiety to its cognate response regulator, which promotes DNA binding and transcriptional control of its target genes. While the regulatory targets and biological functions of many regulatory proteins have been well described (5), a plethora of them remain unexplored.

One of the crucial factors influencing regulatory protein binding to DNA is chromosome topology, determined by chromosome supercoiling and nucleoid-associated proteins (NAPs). The global chromosome topology of bacteria depends on growth phase and environmental conditions, and adjusts transcription in response to both extra- and intracellular conditions. Overall, bacterial chromosome supercoiling is controlled by enzymes called topoisomerases, mainly the TopA-type I topoisomerase, which relaxes DNA (generates positive supercoils), and gyrase, which in contrast introduces negative supercoils (6).

Inhibiting topoisomerase activity or altering topoisomerase levels leads to changes in chromosome topology and affects DNA transactions, including replication and transcription. To date, studies on various bacterial species (*Streptococcus pneumoniae, Haemophilus influenzae, Escherichia coli, Salmonella enterica, Streptomyces coelicolor*) have shown that disturbances in the topological balance affect the transcription of a significant fraction of so-called supercoiling-sensitive genes (7–13).

The binding of NAPs also depends on chromosome topology, with NAPs in turn affecting the binding of other transcription factors (3, 14–18). However, little is known about the cross-talk between chromosome supercoiling and other regulatory systems controlling gene transcription, particularly in response to changes in environmental conditions.

Soil-dwelling bacteria such as *Streptomyces* frequently encounter environmental stress. *Streptomyces* adaptations to the soil environment include their mycelial growth and complex developmental life cycle, which encompasses both spore formation and exploratory growth (19, 20). Vegetatively growing *Streptomyces* cells elongate and branch to generate a network of multicellular hyphae. In response to environmental stimuli, particularly nutrient depletion, sporulation is triggered. Sporulation starts with raising aerial hyphae, within which spore chains subsequently develop. The conversion of multigenomic hyphal cells to chains of unigenomic spores requires chromosome condensation and segregation, accompanied by synchronous septation (19, 21). The progression of the *Streptomyces* life cycle is governed by a set of well-described regulatory proteins (such as those encoded by the *whi* or *bld* genes) (22, 23); however, numerous reports indicate an abundance of less studied regulators and other proteins that also contribute to sporulation regulation (24–29).

*Streptomyces* use a repertoire of biologically active secondary metabolites to thrive in their environmental niche, including numerous antibiotics (approximately 60% of natural antibiotics are *Streptomyces*-produced), immunosuppressants and cytostatics (30). The production of secondary metabolites remains under the control of complex regulatory networks and is coordinated with developmental programmes (31–34). As a free-living organism, *Streptomyces* respond to highly variable conditions using a large number of transcriptional regulators, many of which remain uncharacterized (31). The number of transcription factors encoded by streptomycete genomes ranges from 471 to 1101, and among these, depending on the species, there are 315 to 691 transcriptional regulators and 31 to 76 sigma factors (26). In comparison to other bacterial genera, *Streptomyces* genomes also encode numerous TCSs, the number of which varies depending on the species, ranging from 59 and 117, alongside 13-21 orphan response regulators and 17-39 unpaired/uncharacterized sensor kinases (35–37).

As in other bacteria, chromosome topology plays a critical role in the regulation of gene expression in *Streptomyces*. In contrast to many bacteria, the model *Streptomyces* species *S. coelicolor* possesses only one topoisomerase, the type I TopA, which is essential for viability (38). TopA depletion in *Streptomyces* results in increased DNA supercoiling and altered gene expression, leading to severe growth retardation and sporulation blockage (38, 39). Moreover, disturbances in global DNA supercoiling affect the transcription of up to 7% of *Streptomyces* genes (12). Numerous supercoiling-sensitive genes are grouped into discrete clusters, with one cluster in particular, named SHC (supercoiling hypersensitive cluster), exhibiting extreme DNA supercoiling sensitivity. This region encodes many proteins of unassigned function but also appears to include a two-component system, anti-sigma factors and probable transcriptional regulators. Interestingly, most of the SHC genes are poorly transcribed under standard conditions but are upregulated in response to increased DNA supercoiling (12). Having established that altered DNA supercoiling significantly impacts transcription in *S. coelicolor*, we predicted that altered gene expression may contribute to the sporulation inhibition observed for TopA-depleted strain.

To identify the genes responsible for sporulation and growth inhibition under high supercoiling conditions, we performed random transposon mutagenesis of the TopA-depleted *S. coelicolor* strain, and screened for strains with mutations that suppressed the sporulation blockage associated with high DNA supercoiling. We found that mutations in genes encoding a two-component system named SatKR (SCO3390-89) led to altered transcription of the SHC cluster. We established that the activated response regulator SatR (SCO3389) inhibited growth and sporulation by inducing transcription of SHC genes independently of high DNA supercoiling. Moreover, we confirmed that mutations within SHC prevented the activation of genes within this region and restored growth and sporulation to the TopA-depleted strain. Thus, our results reveal a unique interplay between the two-component system SatKR and chromosome supercoiling in regulating SHC gene expression, with the SHC products subsequently impacting *S. coelicolor* growth and sporulation.

## MATERIALS AND METHODS

### Bacterial strains, plasmids and growth conditions

Basic DNA manipulations were performed according to standard protocols (40). Unless otherwise stated, all enzymes and isolation kits were supplied by Thermo Fisher Scientific (Waltham, MA, US) and NEB (Ipswitch, MA, US). Bacterial media and antibiotics were purchased from DIFCO Laboratories (Detroit, MI, US) and Carl Roth (Karlsruhe, Germany), respectively. The *S. coelicolor* strains used in this study are listed in Supplementary Table 1. Strain construction details are provided in the Supplementary information. The growth conditions, antibiotic concentrations and *S. coelicolor* conjugation procedure were used as described by Kieser et al. (41). To restore TopA levels in the TopA-controlled strain (PS04) (during growth analysis and spore sensitivity assay) the growth medium was supplemented with 1 µg/ml thiostrepton (42). During conjugation with the PS04 strain thiostrepton concentrations of 5 µg/ml were used, unless otherwise stated (it was earlier shown that induction with thiostrepton at concentration higher than 2 µg/ml increase TopA levels only slightly above the wild type).

For growth rate analyses, *S. coelicolor* cultures were inoculated with spores diluted to OD_600_ 0.01/ml in 79 medium. To determine the growth rate, cultures were grown for 48-55 hours in microplates in a final volume of 300 μl using a Bioscreen C instrument (Oy Growth Curves Ab Ltd., Helsinki, Finland), with optical density (OD_600_) measurements being taken at 20-minute intervals. To analyse *Streptomyces* differentiation, strains were plated on solid MS-agar medium and were cultured for 3-7 days.

### Transposon mutagenesis

Random transposon mutagenesis was performed on the TopA-controlled strain (PS04) using the synthetic *Himar1* transposon (3276 bp in length, containing a spectinomycin resistance gene (*aadA*(*1*) and R6Kγ *ori* flanked with ITRs (inverted terminal repeats) (43).

Exconjugants were selected using hygromycin and spectinomycin, in addition to 0.2 µg/ml thiostrepton, to limit transposase induction but increase TopA level. Spores of exconjugants were collected and inoculated into liquid cultures with thiostrepton (0.2 µg/ml); these were cultivated at 39°C overnight to eliminate the pHSM plasmid. The mutant library was then spread for single colonies (to obtain at least 16,000 mutants) on MS-agar plates supplemented with spectinomycin but no thiostrepton (the PS04 strain has a “white phenotype” under these conditions), and grey colonies were screened for. The transposon insertion sites in the selected transposon library clones were identified using a rescue plasmid approach (43). In the MGHM5 strain (PS04 *sco3390*::*Himar1*, *sco2474*::*Himar1*) insertion sites were additionally confirmed by whole-genome sequencing (Genomed, Warsaw, Poland).

### Supercoiling reporter plasmid isolation

The pWHM3Hyg plasmid, which served as a probe of the DNA supercoiling state in vivo, was isolated according to a previously described procedure (42) from *S. coelicolor* strains (MGHM5_RP, MS10 and MS11 – derivatives of analysed mutants modified by pWHM3Hyg introduction), where these strains were cultivated in liquid 79 medium for 24 hours at 30°C. The isolated plasmid DNA was resolved on a 0.8% agarose gel with 2.32 μg/ml chloroquine in TAE buffer. To visualize topoisomers, the gel was stained with ethidium bromide. The topoisomer distribution was analysed using ImageJ Software.

### RNA-Seq and data analysis

For the RNA-seq experiments, RNA was isolated from *S. coelicolor* mycelia obtained from 18-hour cultures in 30 ml YEME/TSB liquid medium. The mycelia were collected by centrifugation, frozen and stored at −70°C for subsequent RNA isolation. RNA was isolated using the procedure described previously by Moody et al. (43), after which the preparations were subjected to digestion with TURBO DNase I (Invitrogen, Waltham, MA, US) and checked using PCR to ensure the samples were free of chromosomal DNA contamination.

Strand-specific cDNA libraries with an average fragment size of 250 bp were constructed, and sequenced using a MiSeq kit (Illumina, San Diego, CA, US) at the Farncombe Metagenomics Facility at McMaster University (Hamilton, Canada). Paired-end 76-bp reads were subsequently mapped against the *S. coelicolor* chromosome using Rockhopper software (44), achieving 1.0-1.5*10^6^ successfully aligned reads per sample. For data visualization, Integrated Genomics Viewer (IGV) software was used (45, 46). The analysis of differentially regulated genes was based on the data generated by Rockhopper software. To calculate the fold change in gene transcription, the normalized gene expression in the control strain (WT or PS04) was divided by the normalized gene expression under particular experimental conditions, delivering information on the fold change, and subsequently, the log2 value of the fold change was calculated. The genes with a q-value (Rockhopper adjusted p-value) greater than or equal to 0.01 and a log2 of the fold change in the range from −1.5 to 1.5 were rejected from the subsequent analysis as not significant. Volcano plots were prepared using R Studio software and the EnhancedVolcano package (R package version 1.10.0, https://github.com/kevinblighe/EnhancedVolcano).

### RT-qPCR

For RT-qPCR analyses, RNA was isolated from *S. coelicolor* mycelia obtained from 24-hour cultures growing in 5 ml liquid 79 medium. Transcription was arrested by adding “STOP solution” (95% EtOH v/v, 5% phenol v/v) (47), and mycelia were harvested by centrifugation and frozen at −80°C. Total RNA was isolated using TRI-Reagent® (Sigma-Aldrich, Saint Louis, MO, US) according to the manufacturer’s procedure. Homogenization was performed in a FastPrep-24™ instrument (MP Biomedicals, Irvine, CA, US) (6 m/s, 2 cycles × 45 s). After centrifugation, RNA was isolated by chloroform extraction, purified on a column (Total Mini RNA, AA Biotechnology, Gdańsk, Poland) and eluted with 50 µl of ultrapure water. The isolated RNA was digested with TURBO DNase (Invitrogen) according to the manufacturer’s instructions at 37°C for 30 minutes. Then, RNA was purified and concentrated using a CleanUp RNA Concentrator (AA Biotechnology) and eluted with 17 µl of ultrapure water. Five hundred nanograms of RNA were used for cDNA synthesis using a Maxima First Strand cDNA Synthesis Kit (Thermo Fisher Scientific) in a final volume of 20 μl. The original manufacturer’s protocol was modified for GC-rich transcripts by increasing the temperature of the first strand synthesis to 65°C and elongation time up to 30 minutes. Subsequently, the obtained cDNA was diluted 5 times and used directly for quantitative PCRs performed with PowerUp SYBR Green Master Mix (Applied Biosystems, Waltham, MA, US). The relative level of a particular transcript was quantified using the comparative ΔΔCt method, and the *hrdB* gene was used as the endogenous control (StepOne Plus real-time PCR system, Applied Biosystems). The sequences of optimized oligonucleotides used in this study are listed in Supplementary Table 2).

### Microscopic analyses

For analysis of spore formation, the tested strains were inoculated at the acute-angled junctions of coverslips inserted at 45° into minimal medium agar plates supplemented with 1% mannitol and cultured for 53 hours to ensure sporulation of all mutant strains. Sporulation of the TopA-controlled (PS04) strain was induced by the addition of 1 µg/ml thiostrepton. Coverslips were fixed with methanol and then mounted using 50% glycerol solution in PBS. DNA was stained with a 2 μg/ml DAPI solution (Molecular Probes, Eugene, OR, US). Microscopic analyses were performed using a Leica microscope (Leica Microsystems, Wetzlar, Germany) with phase contrast imaging. Images were analysed using ImageJ software. The statistical analysis of spore length was performed using R Studio (RStudio Team (2020). RStudio: Integrated Development Environment for R. RStudio, PBC, Boston, MA, US) and the Student’s t-test for paired samples.

### Spore sensitivity assay

To test spore viability, *S. coelicolor* strains were cultured on MS-agar plates for 5 days. Next, the spores were collected and incubated in 5% SDS (sodium dodecyl sulphate) solution for 1.5 hours, washed twice with ultrapure water and resuspended in 0.5 ml of water. Next, serial dilutions were spread on MS agar plates to obtain single colonies.

Subsequently, the number of colonies grown after SDS treatment was compared with the negative control (spores of the same strain collected and incubated in water). Spore viability was calculated as a ratio of the number of colonies obtained for spores treated and untreated with SDS.

## RESULTS

### Screening for suppressors of supercoiling-induced sporulation blockage

TopA is the only type I *S. coelicolor* topoisomerase, and consequently is essential for viability. Its depletion in the TopA-controlled strain (PS04, in which the *topA* gene expression is under the control of the thiostrepton-inducible promoter *tipA,* allowing for an up to 20-fold depletion of TopA levels) leads to increased negative DNA supercoiling (38). Elevated negative DNA supercoiling in turn results in changes in global gene expression and affects the growth rate, sporulation and secondary metabolism of *S. coelicolor* (12, 38). During differentiation of wild-type *S. coelicolor,* white sporogenic (aerial) hyphae mature into chains of grey spores; in contrast, the development of a TopA-depleted strain is inhibited at the aerial hyphal stage, resulting in a “white colony phenotype”. We speculated that inhibition of aerial hyphae maturation may result from changes in the expression of supercoiling-sensitive genes encoding sporulation regulators (12). To identify any such sporulation regulators, we searched for transposon mutations that were able to suppress the TopA depletion phenotype and restore sporulation (grey colonies). To ensure that the transposon insertion frequency was sufficient to cover all 7,825 predicted *S. coelicolor* genes (48), we aimed to obtain a mutant library containing approximately 16,000 clones. Having obtained the representative transposon library (PS04-Tnlib), we searched for mutants that formed grey colonies under TopA-depleted conditions. We identified seven transposants exhibiting this phenotype, and among them, one transposant, termed MGHM5, additionally exhibited a partially restored growth rate upon TopA depletion (with effective depletion being confirmed by Western blotting and RT-qPCR, Fig. S1), both on solid and in liquid media, when compared with its TopA-depleted parental strain (Fig. 1A and 1B). Unlike the TopA-depleted parental strain, which overproduced blue actinorhodin, the TopA-depleted transposon strain did not produce either of the pigmented antibiotics made by *S. coelicolor* (blue actinorhodin or red undecylprodigiosin) (Fig. 1A). Microscopic analysis of spores produced by the TopA-depleted transposon strain confirmed the presence of spore chains, although these were detectable only after prolonged incubation (approximately 53 hours, in comparison to the 48 hours needed to sporulate in the wild-type strain); spore chains could not be detected in the TopA-depleted parental strain (Fig. 1C). Interestingly, the spores produced by the TopA-depleted transposon strain were of varying sizes compared with the wild-type strain and the parental strain in which the TopA level was restored to that of the wild-type strain (PS04, 1 µg/ml thiostrepton). Moreover, spores produced by the TopA-depleted transposon strain were highly sensitive to 5% SDS: only 2.5% survived 1 hour of SDS exposure compared with the 76% spore survival of the same strain with restored TopA levels (Fig. 1D).

**Fig. 1.**
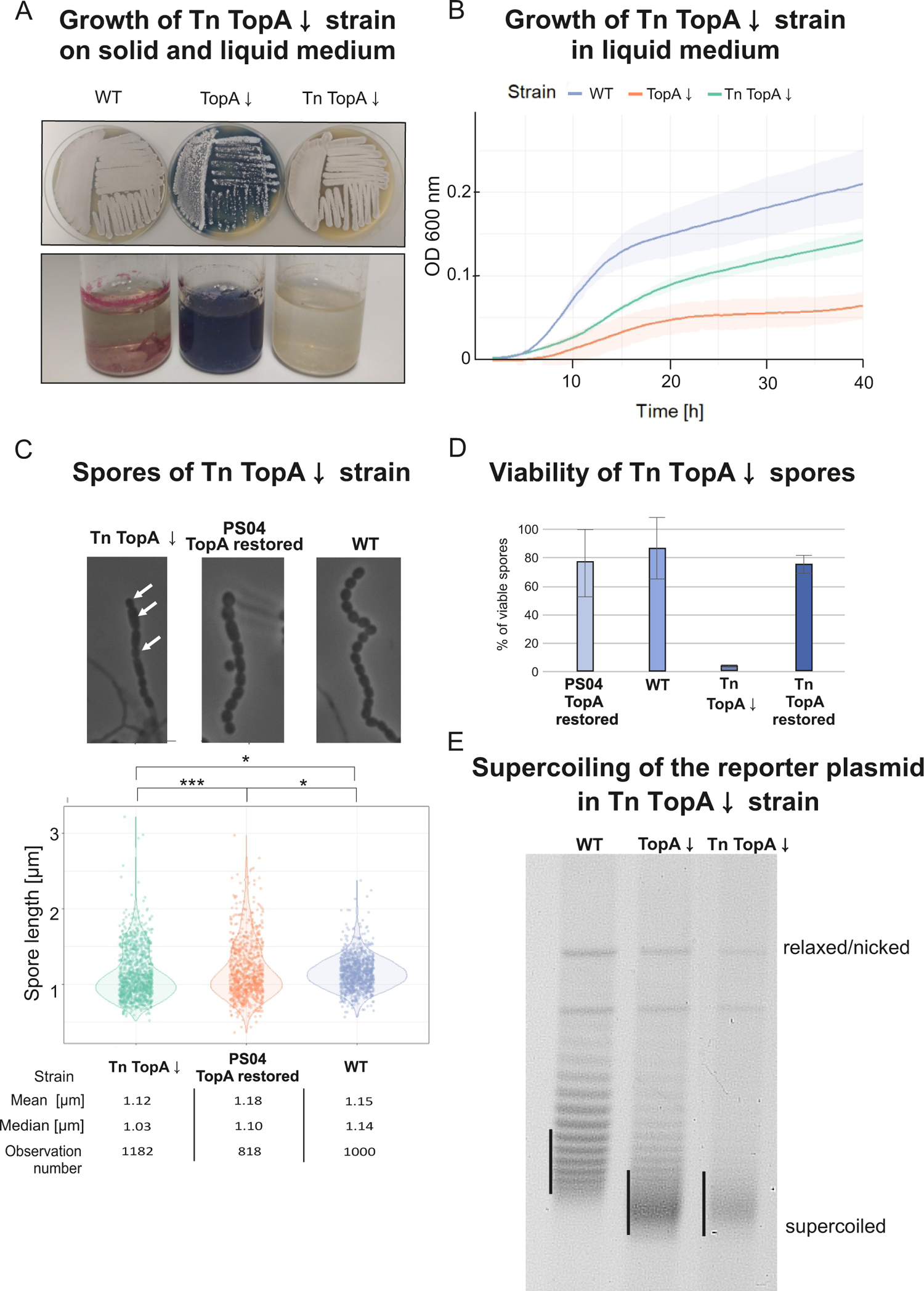
Phenotype of the TopA-depleted transposon strain MGHM5. (Tn TopA↓) **A**. Growth of the TopA-depleted transposon strain (Tn TopA↓) on solid MS agar (upper panel) and antibiotic production in R2 liquid medium (lower panel) compared with wild-type (WT) and TopA-depleted PS04 (TopA↓) strains. The cultures were grown for 72 hours. **B**. Growth curves of the TopA-depleted transposon strain (Tn TopA↓) in liquid 79 medium compared with wild-type (WT) and TopA-depleted PS04 (TopA↓) strain growth. The growth rate was measured in triplicate using a Bioscreen C instrument for 48 hours. **C**. Spores produced by the TopA-depleted transposon strain (Tn TopA↓). Top panel: Phase contrast microscopy images demonstrating representative spore chains of the TopA-depleted transposon strain (Tn TopA↓) and its parental strain PS04 with restored TopA levels (induced with 1 µg/ml thiostrepton) and the wild-type strain (WT) after 53 hours of growth in MM minimal medium (with 1% mannitol). Lower panel: spore size distribution. Asterisks indicate the significance of the p-value (p < 0.05) when comparing mean spore sizes. **D.** Viability of spores of the TopA-depleted transposon strain (Tn TopA↓) after SDS treatment compared with the wild-type strain (WT), PS04 with restored TopA levels and transposon strain (Tn) with TopA level restored. Spores were collected and incubated for 1 hour in 5% SDS at room temperature. The viability percentage was calculated as a ratio of the colony number grown from spores treated and untreated with disrupting agent. **E**. DNA supercoiling of the reporter plasmids pWHM3Hyg or pWHM3Spec isolated from the wild-type strain derivative MS10 (WT), TopA-depleted strain derivative MS11 (TopA↓) and TopA-depleted transposon strain derivative (MGHM5_RP, Tn TopA↓) cultured for 24 hours in liquid 79 medium. The distribution of the reporter plasmid topoisomers was analysed by agarose gel electrophoresis. Black vertical lines indicate the most abundant topoisomers.

Next, we tested whether the increased growth rate and sporulation of the TopA-depleted transposon strain resulted from a restoration of wild type-levels of chromosome supercoiling. To achieve this goal, we isolated a supercoiling reporter plasmid (pWHM3Hyg) from a derivative of the TopA-depleted transposon strain (MGHM5_RP) and established that its negative supercoiling level was similarly high to that of plasmid isolated from the parental TopA-depleted strain derivative, indicating similar chromosome supercoiling (Fig. 1E).

Thus, we successfully identified a transposon mutant in which sporulation and growth rate defects of the parental TopA-depleted strain were restored to wild type levels, and the observed phenotypic effect did not result from restored negative supercoiling.

### Transposon insertion in two-component system-encoding genes influences the expression of a supercoiling-sensitive cluster

We mapped the transposon insertion loci in the MGHM5 strain (by sequencing rescue plasmids and genome sequencing), identifying two transposons: one in *sco3390* and one in *sco2474*. In both cases, the orientation of the *aadA*(*1*) gene within the transposon cassette was the same as that of the disrupted gene. In the first locus (*sco3390*), the transposon cassette was inserted 292 nucleotides downstream of the start codon (Fig. 2). The *sco3390* gene (1206 bp length) was annotated as encoding a putative two-component system kinase, while the genes downstream of this, in a presumable operon, were annotated as encoding a probable cognate response regulator (*sco3389*) and TrmB-like protein (*sco3388*) (49, 50). The second transposition site was located 1020 nucleotides downstream of the start codon of the *sco2474* gene (1644 bp in length), which encodes a putative secreted metalloproteinase (Fig. S2A). Importantly, previously performed RNA-seq analysis (12) showed that while the *sco3388-3390* genes were transcribed during *S. coelicolor* vegetative growth in a supercoiling-insensitive manner (Fig. 3A, left panel), the *sco2474* gene was not expressed during vegetative growth, either in normal or in high DNA supercoiling conditions (i.e. in the TopA-depleted PS04 strain) (Fig. S2B). Moreover, *sco2475,* located downstream of the disrupted gene, was transcribed in the transposon strain, and its expression was not changed due to transposition, suggesting a lack of polar effects associated with this transposon insertion (Fig. S2B). Since *sco2474* was not expressed under any tested conditions, and we knew that transposition in MGHM5 affected not only sporulation but also the vegetative growth rate, we excluded the disruption of *sco2474* as a reason for the restored growth of the TopA-depleted transposon strain and focused our attention on the *sco3390* gene/operon. Transposon insertion in the kinase-encoding *sco3390* gene was expected to affect the level of expression of the downstream response regulator-encoding *sco3389* gene.

**Fig. 2.**
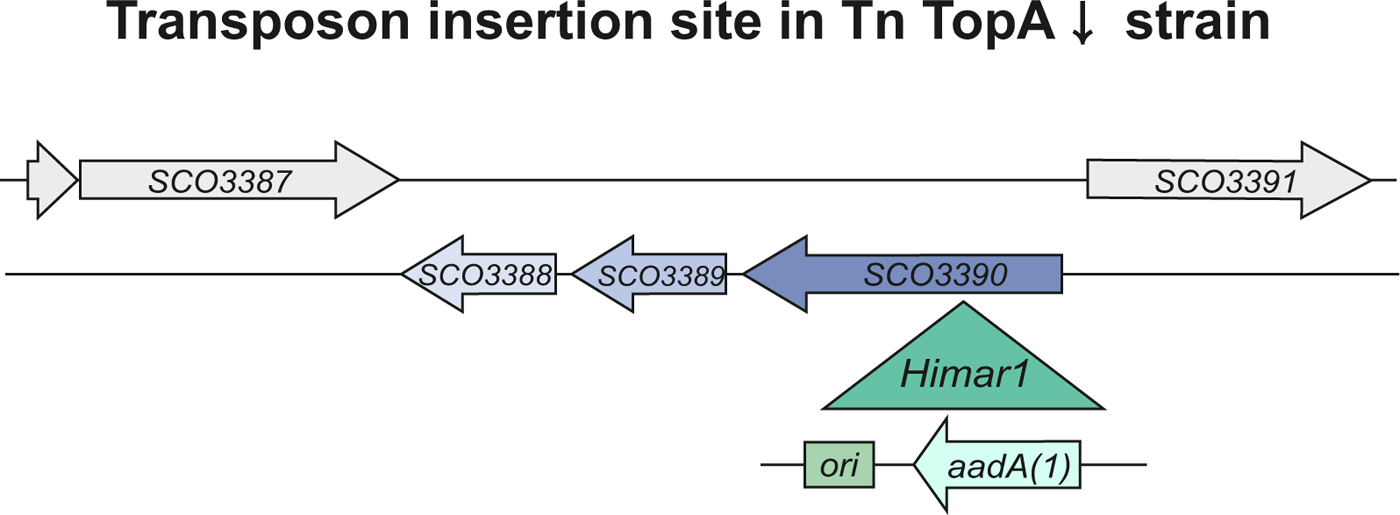
Position of the Himar1 transposon insertion site in the MGHM5 strain. The green triangle shows the insertion site within the *sco3390* gene with the orientation of the inserted *aadA*(*1*) gene. *ori*-origin of replication.

**Fig. 3.**
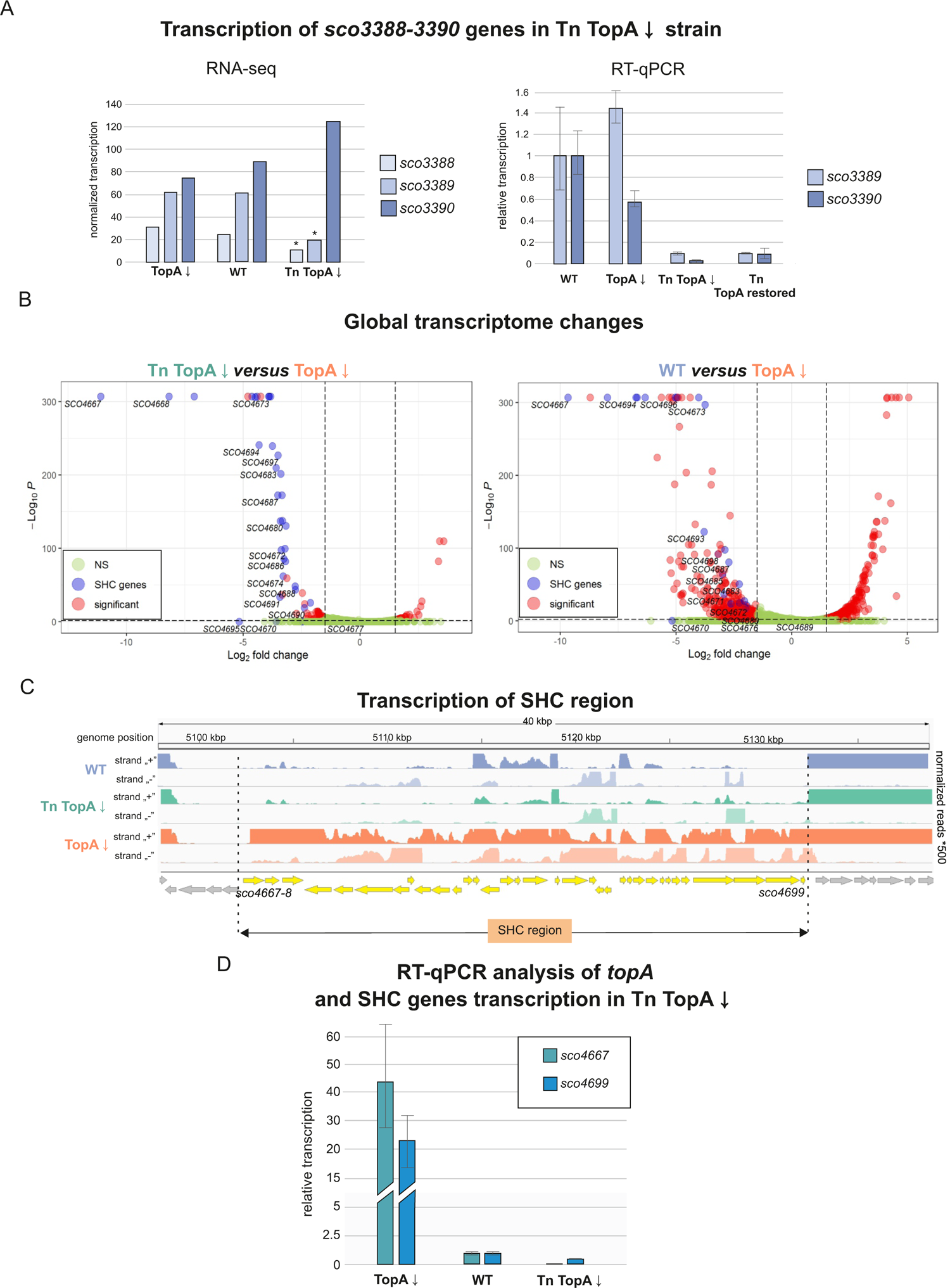
Transcriptional changes in the TopA-depleted transposon mutant MGHM5 (Tn TopA↓) compared with its TopA-depleted parental strain (PS04) and wild-type strain. **A.** Normalized transcription level of *sco3390-sco3388* genes in the TopA depleted transposon strain (Tn TopA↓), transposon strain with TopA level restored (Tn TopA restored), its parental TopA-depleted PS04 strain (TopA↓) and in the wild type (WT), based on RNA-seq (left panel) and the relative transcription level analysed by RT-qPCR (right panel). Asterisks in the RNA-seq analysis indicate statistical significance of the q-value and log2(fold) (q<=0.01 and 1.5<log2(fold)<-1.5). **B**. Volcano plots based on the RNA-seq experiments showing changes in global gene expression in the TopA-depleted transposon strain (Tn TopA↓ strain) compared with its parental TopA-depleted PS04 strain (TopA↓) (left panel), as well as genes affected by supercoiling (changes between the wild-type strain (WT) and TopA-depleted PS04 strain (TopA↓)) (right panel, (12)). Significantly altered transcripts (q<=0.01 and 1.5<log2(fold)<-1.5) are depicted in red, SHC genes are shown in blue, and genes with unchanged transcription are marked in green. **C**. Transcriptional profile of the SHC region in the wild-type strain (blue), TopA-depleted transposon mutant (Tn TopA↓, green) and TopA-depleted parental PS04 strain (TopA↓, orange). SHC genes are depicted in yellow. **D**. Silencing of SHC gene transcription in the TopA-depleted transposon strain (Tn TopA↓) compared with its parental TopA-depleted PS04 strain (TopA↓) and wild-type strain (WT) analysed by RT-qPCR. Primers for amplification of the two genes, the first and last genes of the SHC cluster (*sco4667* and *sco4699*), were used.

Moreover, since the transposon insertion was expected to inactivate the kinase-encoding gene, we further predicted that this mutation would modify the phosphorylation state and activity of its cognate regulator. Transcriptional analysis (RNA-seq, confirmed by RT-qPCR analysis) performed on RNA isolated from liquid cultures of the TopA-depleted MGHM5 transposon strain showed significant *sco3389* downregulation compared with the parental TopA-depleted strain (Fig. 3A and S3). Additionally, we established that the expression of 100 genes changed significantly in the TopA-depleted MGHM5 transposon strain in comparison to its parental TopA-depleted strain. The majority of differentially expressed genes (72 genes) were downregulated in the transposon mutant, with only 28 genes being upregulated in comparison to the parental TopA-depleted strain (Supplementary Table 3 Fig. 3B left panel). Surprisingly, 30 of 72 downregulated genes were concentrated in one region of the chromosome: the supercoiling-hypersensitive cluster (SHC) encompassing 34 genes (*sco4667*-*sco4700*). The majority of the SHC genes (26 of 34 genes) have been previously shown to not be transcribed or transcribed at very low levels under optimal growth conditions but highly induced upon TopA depletion (12) (Fig. 3B right panel and 3C). We confirmed the decreased transcription of SHC genes in the TopA-depleted transposon strain compared with the TopA-depleted parental strain using RT-PCR with primers specific for the first (*sco4667*) and penultimate gene (*sco4699*) of the SHC cluster (Fig. 3D). According to the RT-qPCR results, SHC gene expression was reduced despite TopA depletion, although not to the extent observed in the RNA-seq analysis, which might be due to the different approaches to measure the transcript levels (over the whole gene length vs in a certain position).

In summary, we determined that our TopA-depleted strain carrying a transposon integrated into *sco3390* exhibited an altered transcriptional landscape compared with its parent TopA-depleted strain, with reverted induction of the supercoiling-sensitive SHC genes. This implicated the *sco3389-sco3390-*encoding two-component system in controlling the transcription of the supercoiling-sensitive cluster, and led us to term these gene products SatKR (for **<u>S</u>**HC **<u>a</u>**ctivity controlling **<u>t</u>**wo-component system **<u>K</u>**inase/**<u>R</u>**egulator).

### Changes in SatKR levels affect *Streptomyces* growth and sporulation

To confirm that the SatKR two-component system regulated growth and sporulation in *S. coelicolor* in coordination with chromosome supercoiling, we analysed the phenotypic effects of *satKR* gene deletion and *satR* overexpression in different genetic backgrounds.

Unexpectedly, deleting *satKR* did not affect vegetative growth in liquid medium or differentiation on solid medium in either the wild-type or TopA-depleted backgrounds, (MGM12 and MGP12 strains, respectively) in comparison to parental strains (Fig. 4A).

**Fig. 4.**
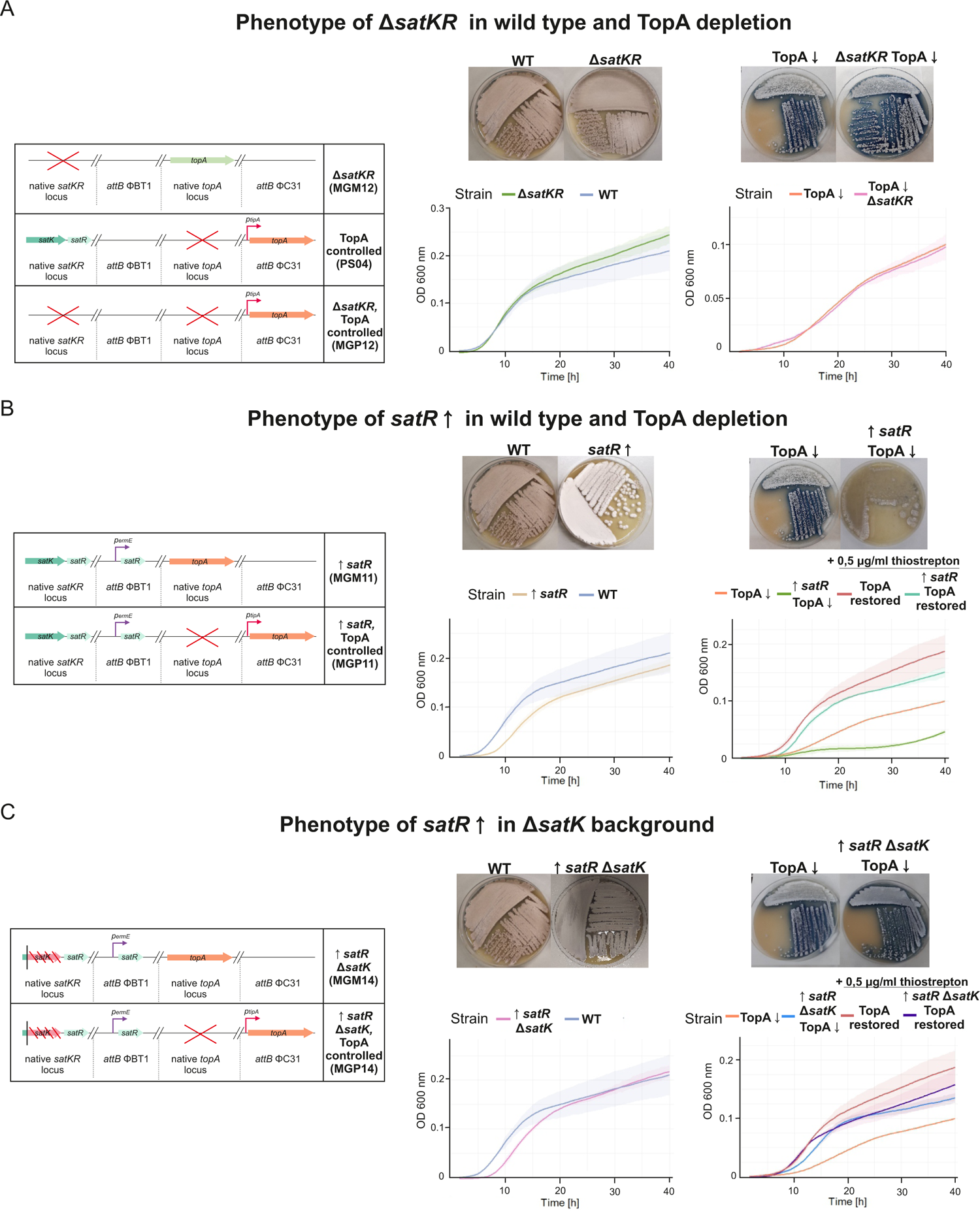
The phenotype of *satKR* mutant strains. **A**. The phenotype of the MGM12 strain with *satKR* deletion in the wild-type background (Δ*satKR*) and the MGP12 strain carrying the *satKR* deletion in the TopA-controlled background (TopA-depleted Δ*satKR* TopA↓and TopA restored with the addition of 0.5 µg/ml thiostrepton inducer: Δ*satKR* TopA restored). On the left: scheme of the mutant strain genotypes. On the right, upper panel: growth and differentiation of strains carrying the Δ*satKR* deletion in the wild-type and TopA-depleted background on MS agar after 48 and 72 hours of growth, respectively. Lower panel: growth curves in liquid 79 broth obtained using a Bioscreen C instrument. Measurements were performed in triplicate every 20 minutes for 42 hours. **B.** The phenotype of strain MGM11 overexpressing *satR* in the wild-type background (*satR*↑) and strain MGP11 overexpressing *satR* in the TopA-controlled background (TopA-depleted: *satR*↑TopA↓ and TopA restored with the addition of 0.5 µg/ml thiostrepton inducer: *satR*↑ TopA-restored). On the left: scheme of the mutant strain genotypes. On the right, upper panel: growth and differentiation of the analysed strains on MS agar after 48 or 72 hours of growth, respectively. Lower panel: growth curves of the analysed strain in liquid 79 broth obtained using a Bioscreen C instrument. Measurements were performed in triplicate every 20 minutes for 42 hours. **C**. The phenotype of the MGM14 strain overexpressing *satR* in the *satK* deletion background (*satR*↑Δ*satK*) and the MGP14 strain overexpressing *satR* in the TopA-controlled *satK* deletion background (TopA depleted: *satR*↑Δ*satK* TopA↓ or TopA restored with the addition of 0.5 µg/ml thiostrepton inducer: *satR*↑Δ*satK* TopA restored). On the left: scheme of the mutant strain genotypes. On the right, upper panel: growth of analysed strains on MS agar for 48 or 72 hours. Lower panel: growth curves of the analysed strain in liquid 79 broth obtained using a Bioscreen C instrument. Measurements were performed in triplicate every 20 minutes for 42 hours.

Sporulation of the Δ*satKR* mutant was also unaffected in the wild-type background, while in the TopA-depleted background, sporulation was not restored, based on plate analyses, and was confirmed microscopically (Fig. S4). Thus, eliminating both components of the SatKR system had different phenotypic effects than the inactivation of the SatK kinase and lowering of the *SatR* transcript levels in transposon strain.

To further explore the role of SatR, we overexpressed *satR* in the presence of its cognate kinase (using an additional copy of the *satR* gene under the control of a strong constitutive *p_ermE_* promoter, with overexpression being confirmed using RNA-seq and RT-qPCR analysis (Fig. S5)) in both the wild-type strain and the TopA-controlled strain (MGM11 and MGP11 strains, respectively, Fig. 4B). Elevated *satR* transcription slightly retarded vegetative growth in liquid cultures and delayed differentiation on solid medium in both the wild-type and the TopA-depleted backgrounds, in comparison to the parental strains (Fig. 4B and S4). Moreover, the *satR*-overexpressing strain with restored TopA levels exhibited somewhat inhibited growth compared with the TopA-restored parental strain (Fig. 4B). These observations suggested that overexpression of *satR* in the presence of the cognate kinase SatK impaired cell growth.

To assess the importance of its cognate kinase on SatR activity, we inactivated *satK* by frame-shift mutation in the wild-type and TopA-controlled backgrounds and overexpressed *satR* in the obtained strains (MGM14 and MGP14 strains, respectively). In contrast to *satR* overexpression in the presence of kinase, elevated *satR* transcript levels in the absence of cognate kinase did not affect the growth rate and development either in liquid or on solid medium (Fig. 4C), compared with the parental strains. These observations suggested that SatK was crucial for SatR activity and its influence on growth and sporulation.

In summary, overexpression of *satR* in the presence of its intact cognate kinase gene (*satK*) led to growth inhibition and delayed sporulation; these effects were lost when *satK* was inactivated.

### The SatKR regulon includes SHC genes

To better understand the biological function of SatKR, we investigated how modulating *satKR* gene expression in different genetic backgrounds affected global gene transcription. Transcriptional analysis showed that deletion of *satKR* in the wild-type background (MGM12) altered the expression of 126 genes (1.5% of all *S. coelicolor* genes) (Supplementary Table 4). Within these genes there were 9 SHC genes (7 of which were supercoiling-sensitive), which we observed to be further downregulated in the *satKR* mutant (despite their very low expression in wild type), indicating that the SatKR system also influenced the expression of the SHC genes under normal supercoiling conditions. Changes in gene expression were also observed when *satKR* was deleted in the TopA-depleted background (strain MGP12), where 67 genes, or 0.76% of all *S. coelicolor* genes, had significantly altered expression (Supplementary Tables 5 and 6). However, the expression of SHC genes activated by high DNA supercoiling in this mutant was unaffected, indicating that the presence of SatR and absence of SatK (as in the transposon mutant) is required for this inhibition.

Next, we established the impact of *satR* overexpression on transcription in the wild-type and TopA-depleted backgrounds (MGM11 and MGP11 strains, respectively). We determined that a high level of SatR in the presence of SatK under normal supercoiling conditions significantly affected the expression of 452 genes (approximately 5% of all genes) (Supplementary Table 7). Among these genes, 351 (nearly 78%) were activated due to *satR* upregulation, while the expression of 101 genes was reduced (Fig. 5A). A similar effect was noted in the TopA depletion background (Supplementary Table 8, 409 genes with altered transcription); however, only 55 genes were equally changed in both wild-type and TopA-depleted conditions. *satR* overexpression in the presence of SatK and normal DNA supercoiling (MGM11) was also found to activate the transcription of 16 genes from the SHC region (Fig. 5B). We confirmed the activation of *sco4667* (first SHC gene) and *sco4699* (penultimate SHC gene) using RT-qPCR (Fig. 5C). In the TopA-depleted background (MGP11 strain), *satR* overexpression did not significantly affect SHC transcription (with one exception being *sco4677*), as under high DNA supercoiling conditions, the transcription of these genes was already highly induced.

**Fig. 5.**
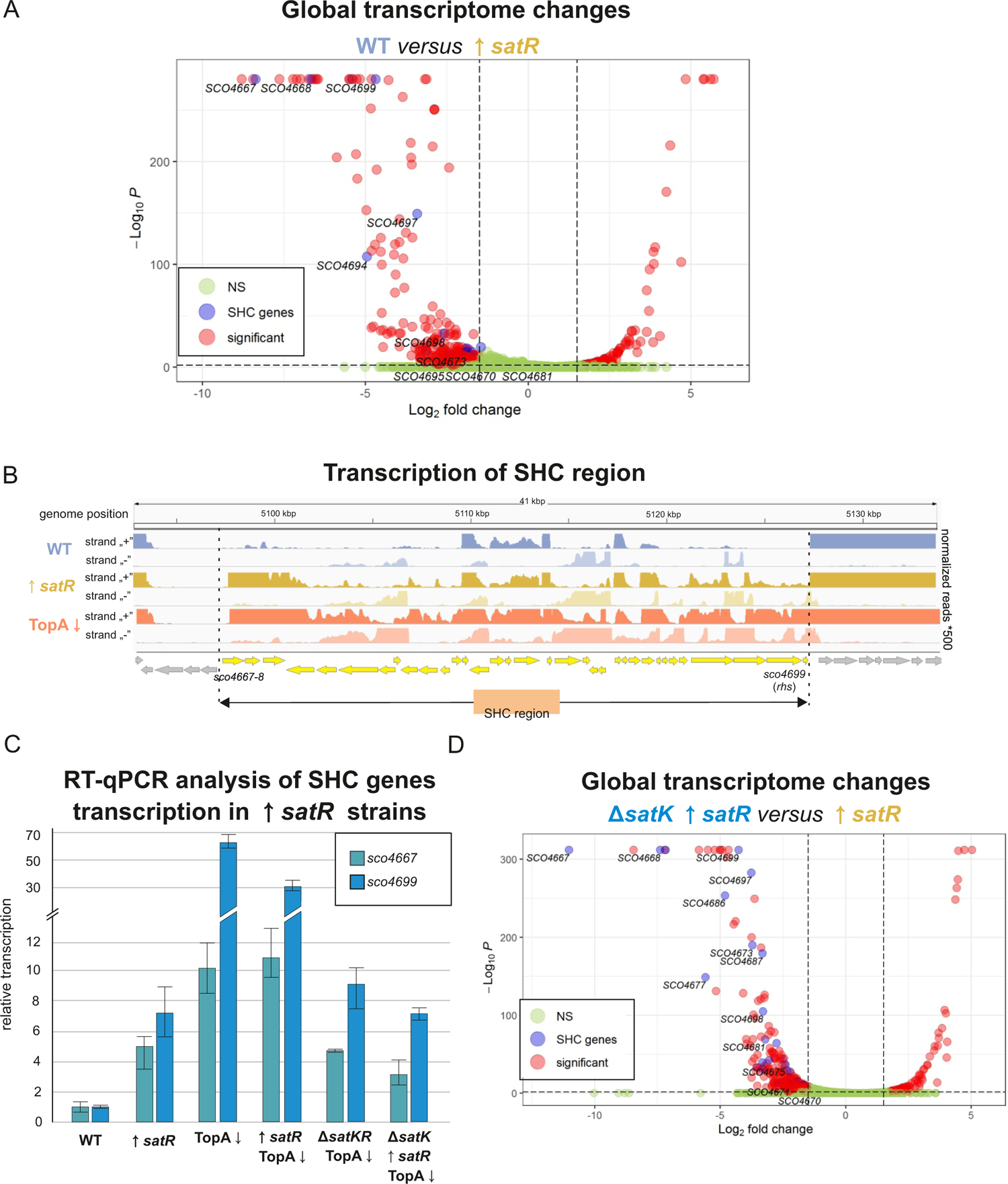
Global transcriptome analysis of strains overexpressing *satR* in the presence of SatK (MGM11) and in the *satK* background (MGM14). A. Volcano plots based on RNA-seq experiments showing changes in global gene expression between the wild-type (WT) and *satR*-overexpressing strain MGM11 (*satR*↑). Significantly altered transcripts (q<=0.01 and 1.5<log2(fold)<-1.5) are depicted in red, SHC genes are shown in blue, and nonsignificant changes are marked in green. **B**. Transcriptional profile of the SHC region in the MGM11 strain overexpressing *satR* (*satR*↑, yellow) compared with its wild-type parental strain (WT, blue) and TopA-depleted PS04 strain (TopA↓, orange). **C.** RT-qPCR analysis of *sco4667* and *sco4699* gene transcription levels in the wild type (WT), TopA-depleted PS04 strain (TopA↓), TopA-depleted MGP12 strain carrying a deletion of *satKR* (Δ*satKR* TopA↓), MGM11 strain overexpressing *satR* in the wild-type background (*satR*↑), MGP11 strain overexpressing *satR* in the TopA-depleted background (*satR*↑ TopA↓) and MGP14 strain overexpressing *satR* in the TopA-depleted *satK* deletion (*satR*↑ Δ*satK* TopA↓) background. **D**. Volcano plots based on RNA-seq experiments showing changes in global gene expression between strain MGM14 overexpressing *satR* in the absence of SatK (*satR*↑ Δ*satK*) and strain MGM11 overexpressing SatR in the wild-type background (*satR*↑). Significantly altered transcripts (q<=0.01 and 1.5<log2(fold)<-1.5) are depicted in red, SHC genes are shown in blue, and nonsignificant changes are marked in green.

While overexpression of SatR in the presence of the SatK kinase activated numerous genes including SHC even under normal supercoiling conditions, its overexpression in the absence of kinase resulted in much less pronounced changes in transcription both in the wild type and increased supercoiling backgrounds (MGM14 and MGP14, respectively) (Supplementary Tables 9 and 10). In conditions of normal supercoiling, overexpression of *satR* in a *satK* mutant background led to transcriptional activation of only 15 genes and repression of 17 genes when compared with wild type. Similarly, overexpressing *satR* in the *satK* mutant, TopA-depleted background led to changes in the expression of 19 genes (14 activated and 5 repressed) compared with the TopA-depleted strain. Furthermore, when comparing the effect of *satR* overexpression in the wild-type and in the *satK* deletion backgrounds (MGP11 and MGP14, respectively), we found that all SHC genes activated by *satR* overexpression in the presence of SatK were silenced when the kinase was absent (Fig. 5D, confirmed by RT-qPCR, Fig. 5C). This evidence strongly suggested that SatK-phosphorylated SatR functions to activate – either directly or indirectly - the SHC genes.

In summary, we confirmed that the SatKR system contributed to the activation of SHC gene transcription. Our transcriptional analyses suggest that SatR functions predominantly as a transcriptional activator, and its activity requires the presence of SatK. Interestingly, our data suggested that both SatKR and supercoiling were sufficient to independently activate SHC transcription. Moreover, the activation of SHC genes either by supercoiling or SatR activity, was correlated with slower growth and sporulation inhibition.

### A single transposon insertion in the SHC induces sporulation under TopA depletion

Our findings suggested that increased SHC expression may be contributing to sporulation defects, given the reduced sporulation observed both in a TopA depletion strain (high supercoiling, high SHC expression) and an *satR* overexpression strain (high SHC expression). Consistent with this possibility was our finding that one of our original transposon mutant strains that restored sporulation to a TopA depleted strain, had an insertion in the penultimate SHC gene *sco4699* (strain MGHM14). *sco4699* encodes a homologue of the Rhs protein from *E. coli*, a secreted toxin that mediates cellular competition (51). In this mutant strain, the *Himar1* transposon was inserted 311 nucleotides downstream of the *sco4699* start codon (Fig. 6A).

**Fig. 6.**
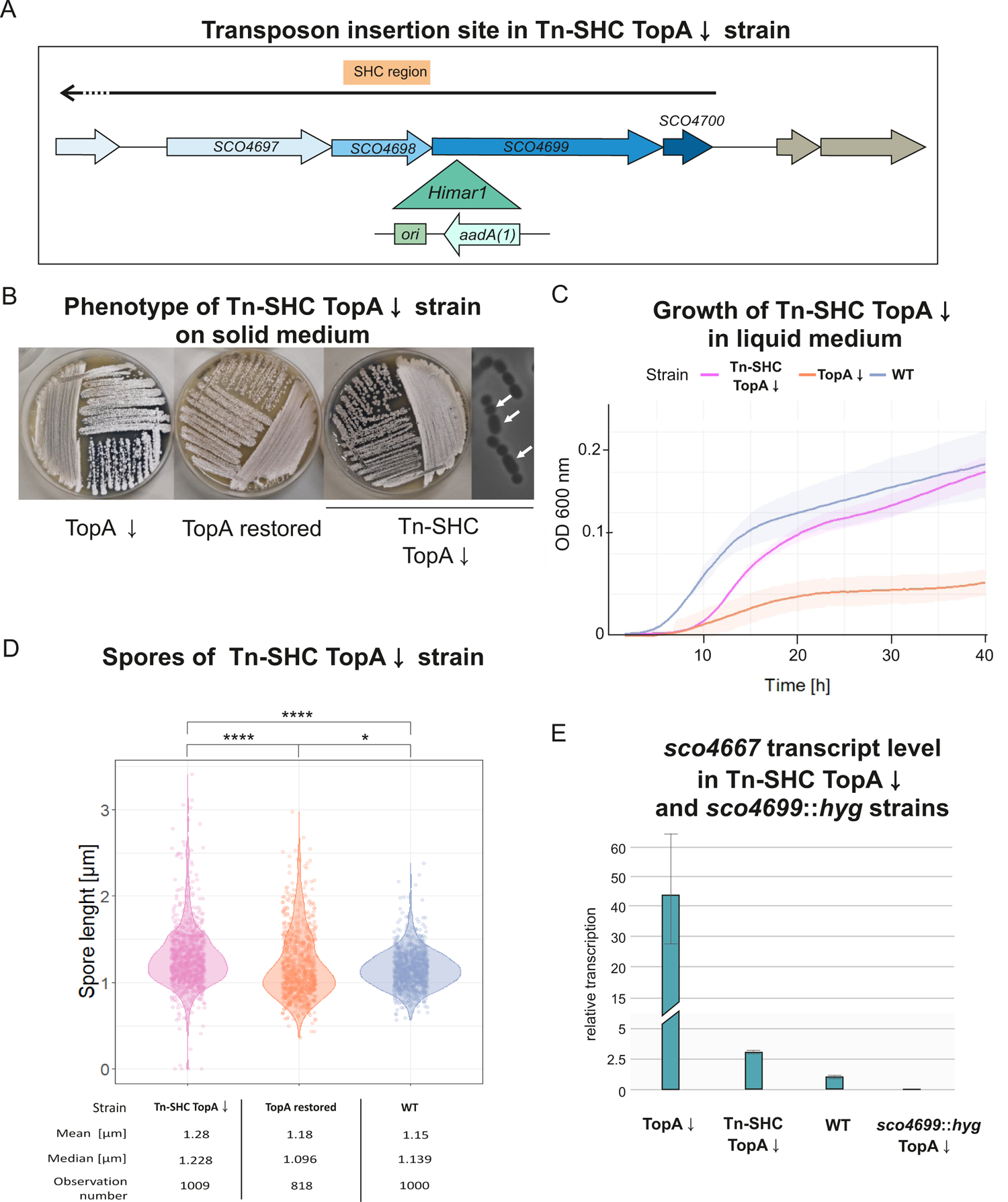
Phenotype of the SHC transposon strain MGHM14. **A.** Himar1 transposon insertion site of the transposon in MGHM14 (Tn-SHC). The green triangle shows the insertion site with the orientation of the inserted *aadA*(*1*) gene. **B**. Growth and differentiation of the TopA-depleted transposon MGHM14 (Tn-SHC TopA↓) compared with the wild-type (WT) and TopA-depleted PS04 strain (TopA↓) on MS agar after 72 hours of growth. **C**. Growth curve of TopA-depleted MGHM14 (Tn-SHC TopA↓) compared with the wild-type (WT) and TopA-depleted strain PS04 (TopA↓) in liquid 79 broth obtained using a Bioscreen C instrument. Measurements were performed in triplicate every 20 minutes for 42 hours. **D**. Spore size distribution in the TopA-depleted MGHM14 strain (Tn-SHC TopA↓), its parental strain PS04 with TopA levels restored by induction with 1 µg/ml thiostrepton (TopA restored) and the wild-type (WT) strain. Asterisks indicate the significance of the p-value (p<0.05) when comparing mean spore sizes. **E**. RT-qPCR analysis of *sco4667* transcription in TopA-depleted MGHM14 (Tn-SHC TopA↓) and TopA-depleted *sco4699* mutant strains (*sco4699*::*hyg* TopA↓) compared with the wild-type (WT) and TopA-depleted PS04 strains (TopA↓).

As was seen for the *satKR*-associated transposon mutant, the growth rate of the TopA-depleted MGHM14 strain (where TopA depletion was confirmed by Western blot analysis, Fig. S6) was partially restored, both on solid and in liquid media, when compared with its TopA-depleted parental strain (Fig. 6B and C). We observed abundant spore chains in the TopA-depleted SHC transposon mutant, while in its parental TopA-depleted strain (PS04), no spore chains could be detected (Fig. 6B). Moreover, similar to what was observed when *satKR* was disrupted by transposon insertion (in the MGHM5 strain), transposon integration into *sco4699* also resulted in the formation of spores of varied sizes (Fig. 6D).

Since *sco4699* product supposedly does not act as transcriptional regulator, we wondered whether the supercoiling-dependent transcriptional regulation of the SHC region might be modified by transposon insertion in *sco4699*, providing a rationale for the restored sporulation. Thus, we used RT-qPCR to test the transcription level of the first SHC gene (*sco4667*) in this mutant background. We found that transposon insertion in *sco4699* silenced expression of the cluster even when TopA was depleted (Fig. 6E), similarly to what had been observed for the *satKR* mutant.

Next, to confirm that the downregulation of the first gene of the SHC resulted from transposon insertion in the last gene of the cluster, we reconstructed the SHC transposon strain by inserting a hygromycin resistance cassette in *sco4699* (at the transposon insertion site) in the TopA-controlled background (MGP20 strain). RT-qPCR analysis confirmed that transcription of the first gene within the SHC region (*sco4667*) was silenced in the TopA-depleted strain (Fig. 6E). This analysis suggested that supercoiling-dependent activation of the SHC genes might be disrupted by insertion in a distant region of the gene cluster.

In summary, the mutation of *sco4699* within the SHC region inhibited cluster activation by supercoiling. Moreover mutations in *sco4699* reinforced the idea that silencing of the SHC under conditions of high supercoiling restores sporulation.

### Deletion of SHC genes (*sco4667-4668*) encoding a two-component system SitKR restores sporulation in conditions of high supercoiling

To further probe the contribution of the SHC genes to the inhibition of sporulation under conditions of high chromosome supercoiling, we sought to test whether specific genes within the SHC region might be responsible. As the first gene of the SHC region (*sco4667*) was annotated as encoding a kinase of another putative two-component system and formed the operon with the *sco4668* gene, encoding its cognate response regulator, we deleted *sco4667-4668* together in the wild-type and TopA controlled backgrounds (MB01 and MB02 strains, respectively).

We found that the *sco4667*-*4668* deletion – like the *satK* transposon insertion and mutation of *sco4699* – partially restored the growth of the TopA-depleted strain in liquid cultures compared with its TopA-depleted parental strain, while having no effect on growth in the wild-type background (Fig. 7A). Sporulation of the *sco4667*-*4668* mutant strain in the TopA-depleted background was also restored; this was confirmed microscopically, but was only observed only after prolonged incubation (120 hours of growth on MS agar) (Fig. 7B). As before, spores of the TopA-depleted *sco4667*-*4668* mutant also exhibited a diversity in size, with a greater mean length than that of wild type (Fig. 7C). Given the ability of this novel two-component system encoded within the SHC to influence growth and sporulation under conditions of high supercoiling, it was named SitKR (**<u>S</u>**porulation **i**nhibiting **<u>t</u>**wo-component system **<u>K</u>**inase and **<u>R</u>**egulator).

**Fig. 7.**
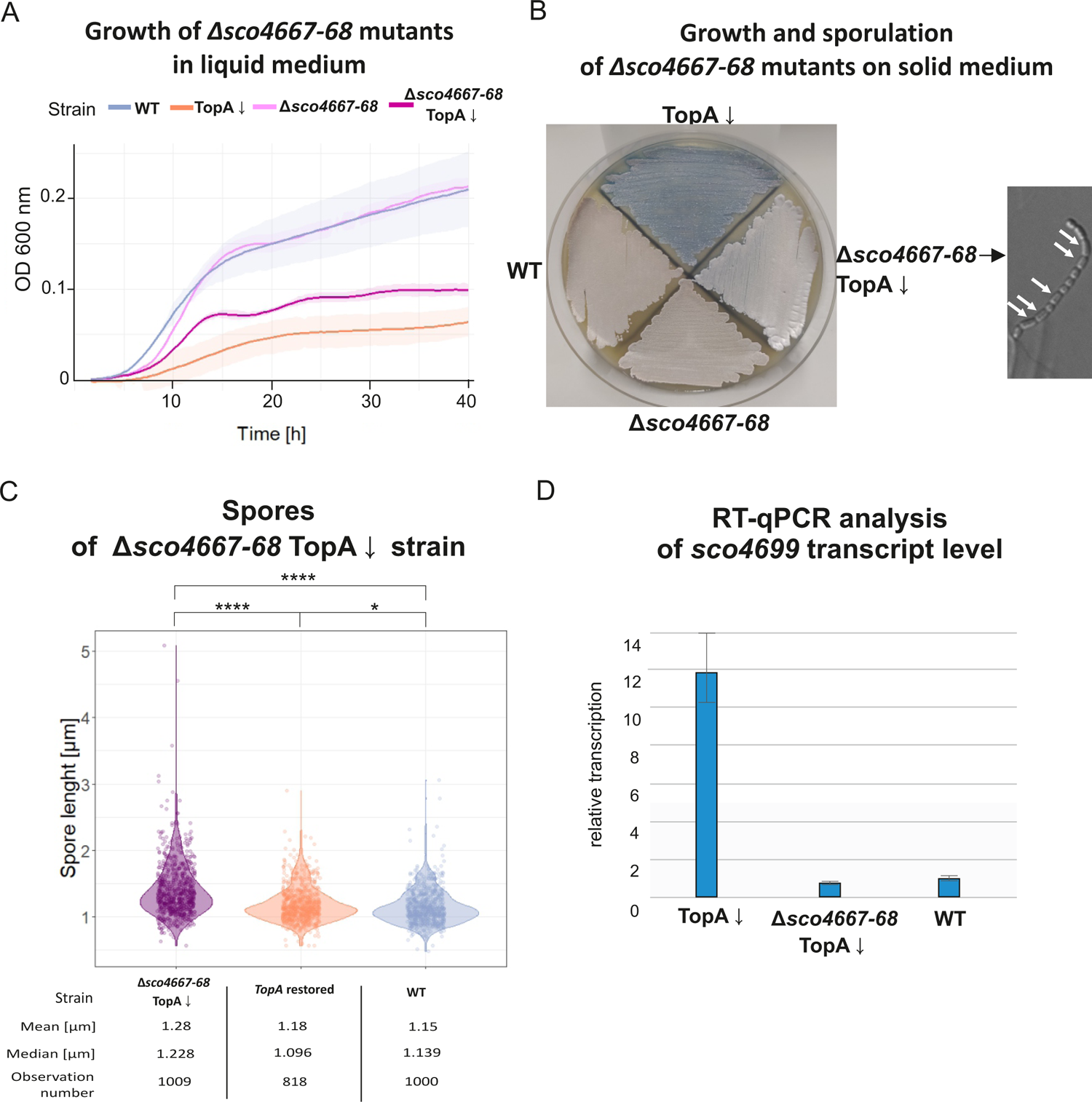
Phenotype of *Δsco4667-4668* mutants in wild-type (MB01) and TopA-depleted backgrounds (MB02). **A**. Growth curves of MB01 (Δ*sco4667-4668*) and the TopA-depleted MB02 strain (Δ*sco4667-4668* TopA↓) compared with the wild-type (WT) and TopA-depleted strain PS04 (TopA↓) in liquid 79 broth obtained using a Bioscreen C instrument. Measurements were performed every 20 minutes for 42 hours. **B**. Growth and differentiation of the MB01 (Δ*sco4667-4668*) and TopA-depleted MB02 (Δ*sco4667-4668* TopA↓) strains, respectively, compared with the wild-type (WT) and TopA-depleted strain PS04 (TopA↓) after 120 hours of growth on MS agar. On the right: microscopic image of the spore chain of the TopA MB02 strain (Δ*sco4667-4668* TopA↓) after 48 hours of growth in MM minimal medium (with 1% mannitol). **C.** Spore size distribution in TopA-depleted MB02 (Δ*sco4667-4668* TopA↓) compared with the parental PS04 strain with restored TopA levels induced with 1 µg/ml thiostrepton (TopA restored) and with wild type (WT). Asterisks indicate the significance of the p-value (p<0.05) when comparing mean spore sizes. **D**. RT-qPCR analysis of the transcription of *sco4699* located at the end of the SHC region in the TopA-depleted MB02 strain (Δ*sco4667-4668* TopA↓) compared with the TopA-depleted PS04 (TopA↓) and wild-type (WT) strain. RNA isolated from a 24-hour culture in liquid 79 broth.

Since we found that insertion in the second-to-last SHC gene abolished TopA depletion-dependent activation of the first SHC gene *sco4667,* we tested whether *sitKR* deletion affected transcription of the penultimate SHC gene. RT-qPCR analysis confirmed that the deletion of *sitKR* genes caused a downregulation of *sco4699* under high supercoiling conditions (Fig. 7D).

This analysis reinforced the idea that mutations of genes within the SHC region can have profound effects on the supercoiling-dependent transcription of more distant genes. Moreover, these results further support the notion that inhibition of SHC gene induction under elevated supercoiling conditions restores the sporulation of TopA-depleted strains. Finally, we have shown that a second two-component system, SitKR, inhibits sporulation when transcriptionally induced by SatKR.

## DISCUSSION

TopA depletion in *Streptomyces* inhibits growth and sporulation. We expected that one of the reasons for this phenotype could be supercoiling-induced changes in the transcription of unknown sporulation/growth regulators. The application of random transposon mutagenesis in the TopA-depleted *S. coelicolor* strain allowed us to identify novel proteins engaged in sporulation regulation. Among the newly identified sporulation regulators was a conserved in *Streptomyces* two-component system named SatKR. The SatKR regulon includes a supercoiling hypersensitive cluster that is also independently activated by TopA depletion. Transposon insertion in the *satK* gene encoding the kinase led to lower levels of SatR, and was observed to revert the TopA depletion-induced activation of SHC and restored growth and sporulation. In contrast, increased SatR levels in the presence of its cognate kinase at wild-type supercoiling levels, led to the activation of transcription of numerous SHC genes, inhibiting *Streptomyces* development.

### The two-component system SatKR controls growth and differentiation by regulating SHC genes

*Streptomyces* two-component systems are known to control antibiotic production, central metabolism, morphology and differentiation (36). Although numerous TCSs in *Streptomyces* affecting these processes have been well-described (*e.g.* CepRS (52), MacRS (53), PhoRP (54, 55), OsdKR (56), AfsQ1/Q2 (57), DraRK (58)), the identification of signals to which TCSs respond has only been successful for a few TCSs (e.g. phosphate concentrations for PhoPR (54), nitrogen concentrations for AfsQ1/Q2 (57) and DraRK (58), iron availability for AbrA1/A2 (59), and redox stress for SenSR (60)). Here, we have established a role for SatKR in regulating growth rate, differentiation and antibiotic production in *S. coelicolor*, but we were unable to predict the SatKR activation signal on the basis of our data. It should be noted that *satKR* genes expression is unresponsive to changes of DNA supercoiling; however, this does not exclude a signal related to changes of DNA topology.

Our results suggested that both the SatR levels in the cell and its activation by the cognate kinase SatK influenced the growth and development of *S. coelicolor*. SatR is a sequence homologue of the DegU response regulator from the *B. subtilis* DegUS two-component system (37.2% identity with 223 aa of overlap). In *B. subtilis,* DegU controls flagellum synthesis, antibiotic production and biofilm formation, inducing the expression of genes involved in matrix formation, competition and nutrient acquisition (61). Interestingly, the *S. coelicolor* genome encodes another DegUS homologue, SCO5784-5785, with a similar percentage identity to DegUS proteins as SatKR. Moreover, SCO5784-5785 has also been shown to influence sporulation as well as antibiotic production; however, its effect on *Streptomyces* differentiation seems to be opposite that of SatKR (62). Markedly, some DegU-regulated genes are induced independently of its phosphorylation (63). Given that SatR stimulates SHC transcription only in the presence of a functional cognate kinase SatK and that reduced levels of SatR in the absence of SatK (in the transposon mutant) lead to the inhibition of SHC transcription despite high DNA supercoiling conditions, we inferred that both SatR states (phosphorylated and unphosphorylated, see below) were able to regulate transcription, although with different effects on the regulated genes. Interesting feature is a localisation of *sco3388* gene downstream *satKR* genes, forming a presumable operon. *sco3388* is similar to *trmB* gene from *B. subtilis*, however, it was previously shown that, in contrast to its homolog, *sco3388* does not determine tunicamycin resistance (64). SCO3388 was indicated to control cell wall integrity and influence spore viability (64), thus its proximity to *satKR* genes may suggest their cooperation in controlling sporulation in *S. coelicolor*.

Since the SatR regulon includes the SHC cluster and SHC is induced in the TopA-depleted strain (12), we concluded that the activation of one or more of the 26 supercoiling-sensitive SHC genes may be responsible for the growth and sporulation inhibition. The SHC region is unique to *S. coelicolor* and shows low synteny amongst *Streptomyces*, even though homologues of individual SHC genes can be found in other species (Fig. S7). The functions of many of the SHC genes remain unknown, but those genes with predicted functions encode two-component systems (*sco4667*-*4668*), two probable regulators (*sco4671* and *sco4673*), a putative Rhs protein (*sco4699*), and the RsfA anti-sigma factor (*sco4677*) (65) that represses SigF, a sporulation-specific sigma factor (66, 67).

The first SHC operon, *sco4667*-*4668,* encoded another novel two-component system named SitKR, which seems to be critical for sporulation and growth. SitKR is conserved among *Streptomyces* species, however its chromosomal location differs among species.

Interestingly, the regulation of the *sitKR* genes expression by SatKR is a phenomenon of a two-component system signalling cascade that has been observed for only few bacterial species, e.g., SsrAB regulation by OmpR/EnvZ and PmrAB by PhoPQ in *Salmonella enterica* (68, 69), and RseDE by PhoPR in *B. subtilis* (70). *sitKR* genes are among the most significantly induced by TopA depletion and *satR* overexpression. Whereas deletion of *satKR* did not restore sporulation in the TopA-depleted strain, a strain in which the *sitKR* genes had been deleted was capable of producing spores. The regulon of *sitKR* has not been established; however, we found that operon deletion abolished supercoiling induction of other genes in the SHC (e.g. *sco4699*). While SitR may directly control the expression of *sco4699* (and possibly other SHC genes), there may also be synchronized regulation between (at least) the first (*sitKR*) and last genes (*sco4699-4700)* within the SHC region. We cannot exclude that upregulation of *sco4677* (encoding anti-sigF) by SatR (and possibly by SitKR) or by increased DNA supercoiling can significantly contribute to sporulation inhibition.

The significance of SHC gene induction for the inhibition of growth and sporulation of the TopA-depleted strain was reinforced by the identification of transposon mutations in the next to the last gene of the SHC region (*sco4699*), which led to restored sporulation in the TopA-depleted strain. *sco4699* encodes a homologue of the Rhs protein from *E. coli*, where Rhs is a secreted toxin that mediates cellular competition (51). Importantly, transposon insertion in *sco4699* also abolished the expression of the first SHC gene, *sitK*. It is possible that the observed effect on sporulation and growth in the SHC transposon strain was due to Rhs inactivation, but we consider the abolished induction of *sitKR* by TopA depletion to be a more likely explanation for the restored sporulation in the transposon strain.

Thus, abolished induction of the SHC region upon TopA depletion in three distinct mutant strains (transposon in *satK*, transposon in *rhs*, and deletion of *sitKR*) restored growth and sporulation despite lowered TopA levels and high chromosomal supercoiling. Notably, spores produced by all the strains with high DNA supercoiling and inhibited SHC transcription had varied sizes and impaired resistance to damaging agents. This phenomenon could result from a lack of additional factors that are needed for proper spore formation and maturation, which may be encoded outside the SHC region. Nevertheless, we established that genes within the SHC region were under the control of the SatKR two-component system and included sporulation and growth rate inhibitors. We speculate that in response to unidentified factors, SatKR induces SHC genes to support slower growth and inhibit sporulation under unfavourable, possibly DNA topology-affecting conditions. Given that a similar induction of SHC genes results from increased supercoiling, we hypothesize that both factors, i.e., changes in chromosome supercoiling and SatKR activation, would independently prevent sporulation under specific environmental conditions, possibly connected with chromosome damage.

### The two-component system SatKR collaborates with chromosome supercoiling in the regulation of gene expression

SHC genes are subject to no or low transcription in the wild-type *S. coelicolor* strain under standard culture conditions, but they are activated by high DNA supercoiling as a result of TopA depletion. However, the presumed low levels of SatR resulting from kinase inactivation (transposon mutant, with low *satR* transcript levels) completely abolished SHC induction by high DNA supercoiling. Importantly, the deletion of *satKR* genes together did not abolish SHC activation by increased DNA supercoiling. Thus, we suggest that in addition to the presence of the kinase, the SatR phosphorylation state may also depend on SatR protein level. Taking into account the possibility of nonspecific activation of SatR by other cellular factors (e.g., acetyl phosphate (AcP) (71) or non-cognate TCS kinases (72)), we suggest that SatR overproduction results in inefficient or nonspecific modification of SatR. Additionally, considering the potential phosphatase activity associated with many kinases, at this stage we cannot predict the phosphorylation state of SatR in presence and in absence of SatK; however we assume that it is different in those two genetic backgrounds. Since we established that deletion of *satKR* genes had different effects on growth and gene expression than a transposon inserted in *satK*, we speculate that upon *satK* inactivation, SatR in its inactive state may also bind to DNA and control the expression of SHC genes, reversing their activation by high supercoiling. This could be achieved either by different DNA binding specificities/affinities for phosphorylated and unphosphorylated SatR or different abilities to form higher order complexes or interact with RNA polymerase. Our speculation here is supported by evidence showing that other two-component system response regulators can act when unphosphorylated (73), e.g., OmpR (74, 75), PhoP (76), SsrB (77) and DegU (63).

Possible explanations for the coordinated regulation of gene transcription by DNA supercoiling and transcription factor activity could include the formation of a supercoiled DNA loop due to transcription factor binding, or supercoiling-dependent binding of transcription factors. Synchronized activation of the SHC region could suggest formation of the loop encompassing this region, stabilized by SatR. Moreover, loop formation could depend on global DNA supercoiling. There is published evidence for DNA looping by some repressor proteins promoted by supercoiling, e.g., LacI in *E. coli* or bacteriophage λ (78, 79), because DNA compaction increases the likelihood of bringing together distant binding sites. Therefore, the affinity of SatR for DNA could be altered not only by its phosphorylation but also by DNA supercoiling. Indeed, the binding of some response regulators has been shown to be influenced by DNA supercoiling, e.g., OmpR binding in *Salmonella enterica* (80).

Elevated *satR* transcription in the presence of kinase activated SHC transcription in the wild-type strain, showing that the SatKR system could control the SHC region independently of DNA supercoiling. In other bacteria species, it has also been suggested that two-component systems might influence the DNA supercoiling state by interacting with topoisomerases or altering their activity. In *Staphylococcus aureus,* deletion of the ArlSR two-component system elevates DNA supercoiling (81), while the *E. coli* RstB response regulator interacts with TopA and increases its activity (82). Although we observed elevated supercoiling in the TopA-depleted strain with a transposon insertion in *satK,* and lowered *satR* transcript levels, we cannot exclude the possibility that local DNA topology changes when SatR binds.

We propose a model in which SatR and DNA supercoiling (caused by TopA depletion), both independently and cooperatively, regulate SHC transcription (Fig. 8). SatR, as a two-component system regulator in the presence of SatK kinase, activates SHC transcription.

**Fig. 8.**
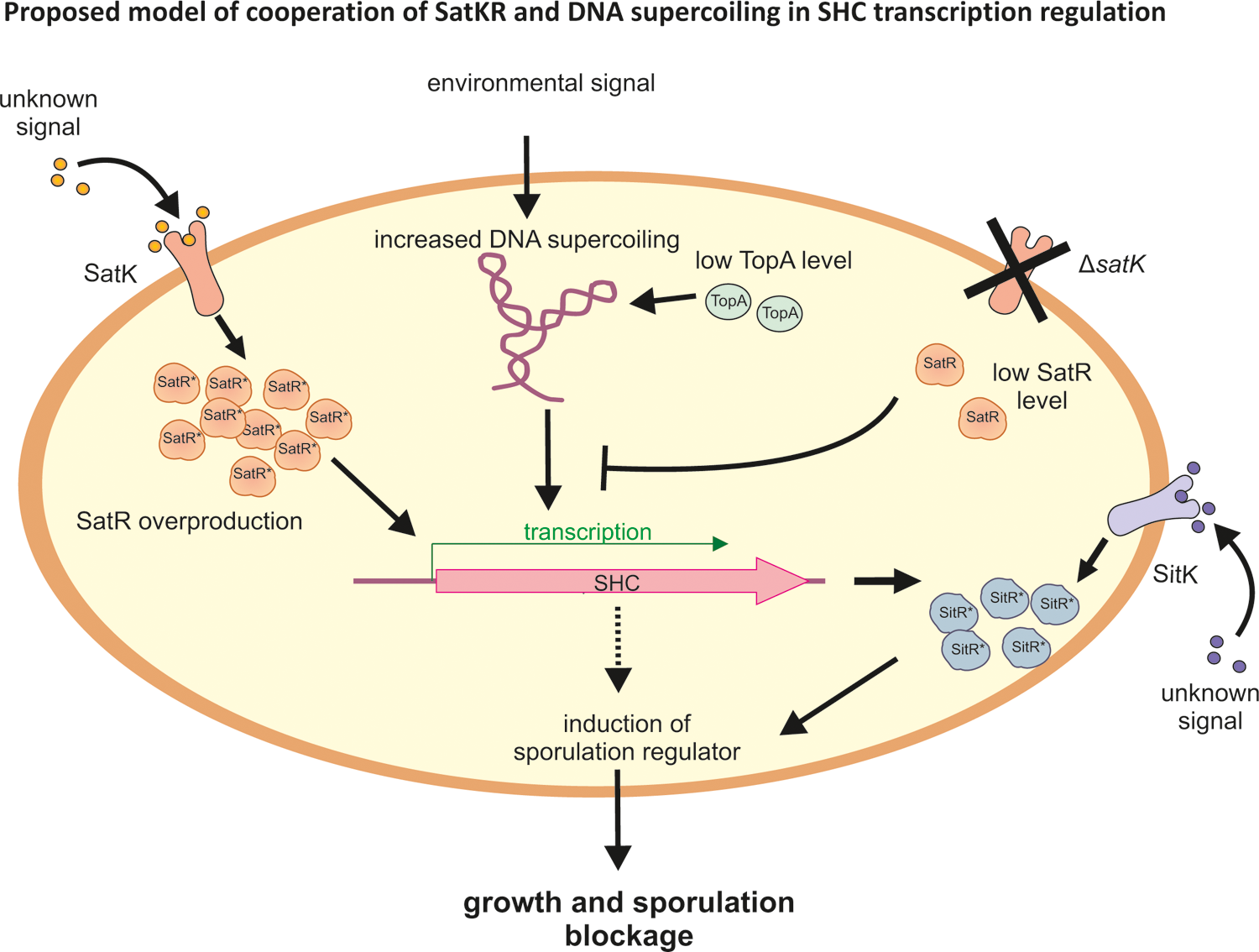
Proposed model of cooperation of SatKR, SitKR and DNA supercoiling in SHC transcription regulation and sporulation inhibition. SHC genes are activated by elevated DNA supercoiling. In the absence of SatK and at low level, SatR inhibits supercoiling-dependent activation of SHC transcription. At high intracellular concentration and when activated by the cognate kinase SatK, SatR induces transcription of SHC genes, independently of supercoiling. Among SHC products there is SitKR two-component system, which is involved in sporulation inhibition. Contribution of other SHC encoded proteins in the sporulation inhibition is also probable.

However, in the absence of kinase, SatR also regulates the transcription of SHC genes, reversing their activation by high supercoiling. Induction of SitKR, an SHC-encoded two-component system, inhibits sporulation; however, it is still unclear if this sporulation defect is direct effect or if other SHC-encoded proteins are involved. Our results are indicating that there is an unprecedented complex regulatory interplay that connects DNA supercoiling with a cascade of two-component regulatory systems, in controlling growth and sporulation in *Streptomyces*.

## DATA AVAILABILITY

RNA-seq data are available at ArrayExpress (EMBL-EBI); accession number E-MTAB-10835.

## ACKNOWLEDGMENTS

We thank Dr Govind Chandra (John Innes Centre, Norwich, UK) for the bioinformatics analysis of SHC gene synteny among *Streptomyces*. This work was supported by the Polish National Science Centre (OPUS grant number 2014/15/B/NZ2/01067 to D.J.); Natural Sciences and Engineering Research Council of Canada (Discovery grant number RGPIN-2020-07197 to M.E.) and Canadian Institutes of Health Research (Project grant number PJT-162340 to M.E.).

## Supplementary figures

**Fig. S1. Analysis of TopA level in the transposon MGHM5 strain. A.** Western blot analysis of TopA level in the TopA-depleted MGHM5 (Tn TopA↓) and TopA-restored (Tn TopA restored) transposon strains compared with the wild-type (WT) strain. **B.** RT-qPCR analysis of *topA* transcript level in TopA-depleted MGHM5 (Tn TopA↓) compared with the TopA-depleted parental PS04 strain (TopA↓) and wild type (WT).

**Fig. S2. The second transposon insertion site in transposon strain MGHM5 A.** Schematic position of the second transposon insertion site within the *sco2474* gene. The green triangle shows the Himar1 insertion site with the orientation of the inserted *aadA*(*1*) gene. **B.** Transcriptional profile of *sco2472-2475* genes in the wild-type strain (WT, blue), TopA-depleted transposon mutant (Tn TopA↓, green) and TopA-depleted parental PS04 strain (TopA↓, orange).

**Fig. S3.** Transcriptional profile of *sco3388*-*3390* genes in the TopA-depleted transposon MGHM5 strain (Tn TopA↓, green) compared with its parental TopA-depleted PS04 strain (TopA↓, orange) and wild type (WT, blue). The green triangle shows the insertion site of the Himar1 transposon within the *sco3390* gene. Black arrows show the localization of *sco3390* RT-qPCR primer binding sites.

**Fig. S4.** Microscopy images of differentiating cultures of the wild-type (WT), TopA-depleted (TopA↓), *satKR* mutant and *satR-*overexpressing strains in the wild-type (MGM12 and MGM11) and TopA-depleted (MGP12 and MGP11) backgrounds. Images were obtained after 52 hours of growth on MM medium supplemented with 1% mannitol under a Leica phase-contrast microscope.

**Fig. S5. Analysis of *satR* transcription. A**. Normalized expression level of *satR* in RNA-seq experiments in the *satR*-overexpressing strain MGM11 (*satR*↑) and TopA-depleted *satR*-overexpressing strain MGP11 (↑*satR* TopA↓) compared with the parental wild-type (WT) and TopA-depleted PS04 (TopA↓) strains (12). The experiment was performed using 18-hour cultures in YEME-TSB broth. **B**. Relative transcription level of *satR* in the RT-qPCR experiment, performed using 24-hour cultures in 79 liquid medium of the *satR* overexpressing strain MGM11 (*satR*↑) and TopA-depleted *satR* overexpressing strain MGP11 (↑ *satR* TopA↓) compared with the parental wild-type (WT) and TopA depleted PS04 (TopA↓) strains, respectively.

**Fig. S6 Analysis of TopA level in the transposon MGHM14 (Tn-SHC) strain.** Western blot analysis of TopA level in the TopA-depleted MGHM14 (Tn-SHC TopA↓) and TopA-restored (Tn TopA restored) transposon strains compared with the wild-type (WT) strain.

**Fig. S7. Synteny of the SHC genes among different *Streptomyces* species.** The SHC genes (*sco4667* to *sco4700*) were reciprocally blasted against 60 *Streptomyces* genome (including *S. coelicolor*) reference sequences from GenBank (full list of genes in Supplementary Table 11). The remaining 42 *Streptomyces* reference genomes do not encode any genes annotated within SHC. SHC genes activated in response to elevated supercoiling are depicted in orange, and nonsensitivity to topology changes is depicted in green.

